# BECN1 F121A mutation increases autophagic flux in aged mice and improves aging phenotypes in an organ-dependent manner

**DOI:** 10.1101/2022.01.12.475960

**Authors:** Salwa Sebti, Zhongju Zou, Michael U. Shiloh

## Abstract

Autophagy is necessary for lifespan extension in multiple model organisms and autophagy dysfunction impacts age-related phenotypes and diseases. Introduction of an F121A mutation into the essential autophagy protein BECN1 constitutively increases basal autophagy in young mice and reduces cardiac and renal age-related changes in longer-lived *Becn1^F121A^* mutant mice. However, both autophagic and lysosomal activity have been described to decline with age. Thus, whether autophagic flux is maintained during aging and whether it is enhanced in *Becn1^F121A^* mice is unknown. Here we demonstrate that old wild type mice maintained functional autophagic flux in heart, kidney and skeletal muscle but not liver, and old *Becn1^F121A^* mice had increased autophagic flux in those same organs compared to wild type. In parallel, *Becn1^F121A^* mice were not protected against age-associated hepatic phenotypes but demonstrated reduced skeletal muscle fiber atrophy. These findings identify an organ-specific role for the ability of autophagy to impact organ aging phenotypes.

## Introduction

Autophagy is an evolutionary conserved lysosomal degradative process essential for the maintenance of cellular homeostasis and the promotion of cell survival under stress conditions. As a result, dysregulation or diminished autophagy activity impacts a wide variety of diseases, including age-related diseases, as well as in aging^1,2^. Indeed, multiple lines of evidence over the past 30 years have demonstrated that autophagy activity declines with age in diverse organisms^3^. Lysosomal protease activity is reduced in aged *C. elegans*^4^ and defective lysosomes and autophagic vacuoles accumulate with age in mouse liver ^5,6^. Moreover, the expression of several macroautophagy/autophagy genes decreases over time in Drosophila^7–9^. Likewise in mammals, levels of the essential autophagy proteins LC3, ATG5 and ATG7 decline with age in mouse brain as well as both muscle and human muscle^10,11^. In addition to declining during normal aging, expression of ATG proteins is also reduced in the setting of age-related diseases such as cardiomyopathy, neurodegenerative diseases and osteoarthritis^12,13^. Genetic studies in organismal models provided more direct evidence of the importance of autophagy in longevity and confirmed the original finding that knock-down of the autophagy gene *bec-1* (encoding BECN1) abrogates the lifespan extension of long-lived mutant worms^14,15^. Similarly, loss-of-function genetic studies indicate that autophagy is essential for lifespan extension in long-lived flies^3^ and reciprocally, induction of autophagy by the overexpression of ATG8 and ATG1 increases lifespan in flies ^7,16^. Since deletion of autophagy genes results in neonatal lethality in mice^17^ and systemic deletion of autophagy genes in adult mice results in early lethality ^18^, genetic studies of autophagy and aging in mammals have taken advantage of mice with tissue-specific deletion of *Atg* genes^11,19–23^. In these models, decreased autophagy results in multiple defects including the accumulation of dysfunctional organelles and protein aggregates that are also found in aging tissues of otherwise non-genetically modified animals ^24^.

The mammalian protein BECN1, orthologue of the yeast protein Atg6, is an essential component of the class III phosphatidylinositol 3-kinase (PtdIns3K) complex that promotes initiation of autophagosome formation. The function of BECN1 in autophagy is negatively regulated by binding to BCL2. Disruption of the BECN1:BCL2 complex enhances the lipid kinase activity of the BECN1-PtdIns3K complex and subsequent induction of autophagy ^25–27^. Transgenic mice bearing a *Becn1*^F121A^ knock-in (KI) mutation that disrupts BECN1 binding to BCL2 represent a unique autophagy gain-of-function mouse model that provided genetic evidence that mice with constitutively increased autophagy have an extended lifespan and improved cardiac and renal aging ^28,29^. In addition, constitutively increased autophagy in *Becn1*^F121A^ KI mice also prevented the age-related decline in neurogenesis and olfaction^30^. Whether autophagy declines in all mouse tissues equally during aging and whether increased autophagy could alleviate the age-associated dysfunction of all tissue/organ during aging is unknown. Moreover, a recent study monitoring autophagy in *C. elegans* during aging demonstrated that autophagic flux is reduced with age in all tissues/organs but lifespan extension by different interventions relies on autophagy in a tissue-specific manner^31^. Thus, whereas increased autophagy has been demonstrated to improve mouse lifespan and healthspan at a whole-body level, the tissue-specific regulation of autophagy during mammalian aging remains to be explored.

In this study, we investigated whether age-related phenotypes could be improved in liver and skeletal muscle of longer-lived *Becn1*^F121A^ KI mice and determined the impact of BECN1^F121A^ mutation on autophagic flux in the liver, heart, kidney and skeletal muscle of old mice.

## Results

We previously reported that the *Becn1*^F121A^ knock-in (KI) homozygous mice that have a constitutive increase of basal autophagy have extended lifespan and delayed age-related cardiac and renal pathological phenotypes^28^. To further investigate if the KI mutation could delay aging phenotypes of other organs, we measured lipid accumulation and fibrosis in aged KI and control (WT) mice. Using 20 month-old WT and KI mice, we analyzed H&E stained tissue sections (Fig. 1A) and quantified the percentage of liver area covered by lipid droplets. Contrary to our expectations, there was no statistically significant difference in lipid accumulation between old KI and WT mice, despite a slight trend towards a greater number of KI mice with less lipid content (Fig. 1B). Next, we evaluated age-related hepatic fibrosis on sections stained with Masson’s trichrome (Fig. 1C and 1D) and similarly, did not observe a statistically significant difference in liver fibrosis between old KI and WT mice. These results indicated that the KI mutation did not protect mice from age-related hepatic pathological changes and led us to ask if the level of autophagy was still higher in the livers of old KI mice compared to old WT mice. Indeed, we previously demonstrated that the KI mutation increased basal autophagy in mouse livers in 6 month-old young adult mice though it has been well characterized that hepatic autophagy declines with age^28,32^. To monitor autophagy in old mice, we studied KI and WT mice that had been crossed to transgenic mice expressing GFP tagged LC3 ^33,34^. In the livers of old mice, we observed no difference in the number of GFP-LC3 puncta between KI and WT mice (Fig. 1E and F1). To assess autophagic flux, we treated mice with the autophagy inhibitor chloroquine (CQ). In CQ treated old mice, neither KI nor WT mice had a statistically significant increase in accumulation of GFP-LC3 puncta in the livers compared to untreated mice, and similarly, we did not observe a significant difference in flux when comparing the livers of old KI and WT mice. There was also no change in the protein expression of autophagy receptor and substrate SQTM1/p62 in the livers of old KI and WT mice as evaluated by western blot, thus confirming similar autophagic flux between old KI and WT mice in the liver (Fig. 1G). Taken together, these results demonstrate that in contrast to the livers of young mice, old *Becn1*^F121A^ KI mice did not display increased liver autophagy, which correlated with the inability of the KI mutation to impact liver aging phenotypes.

**Figure 1.**
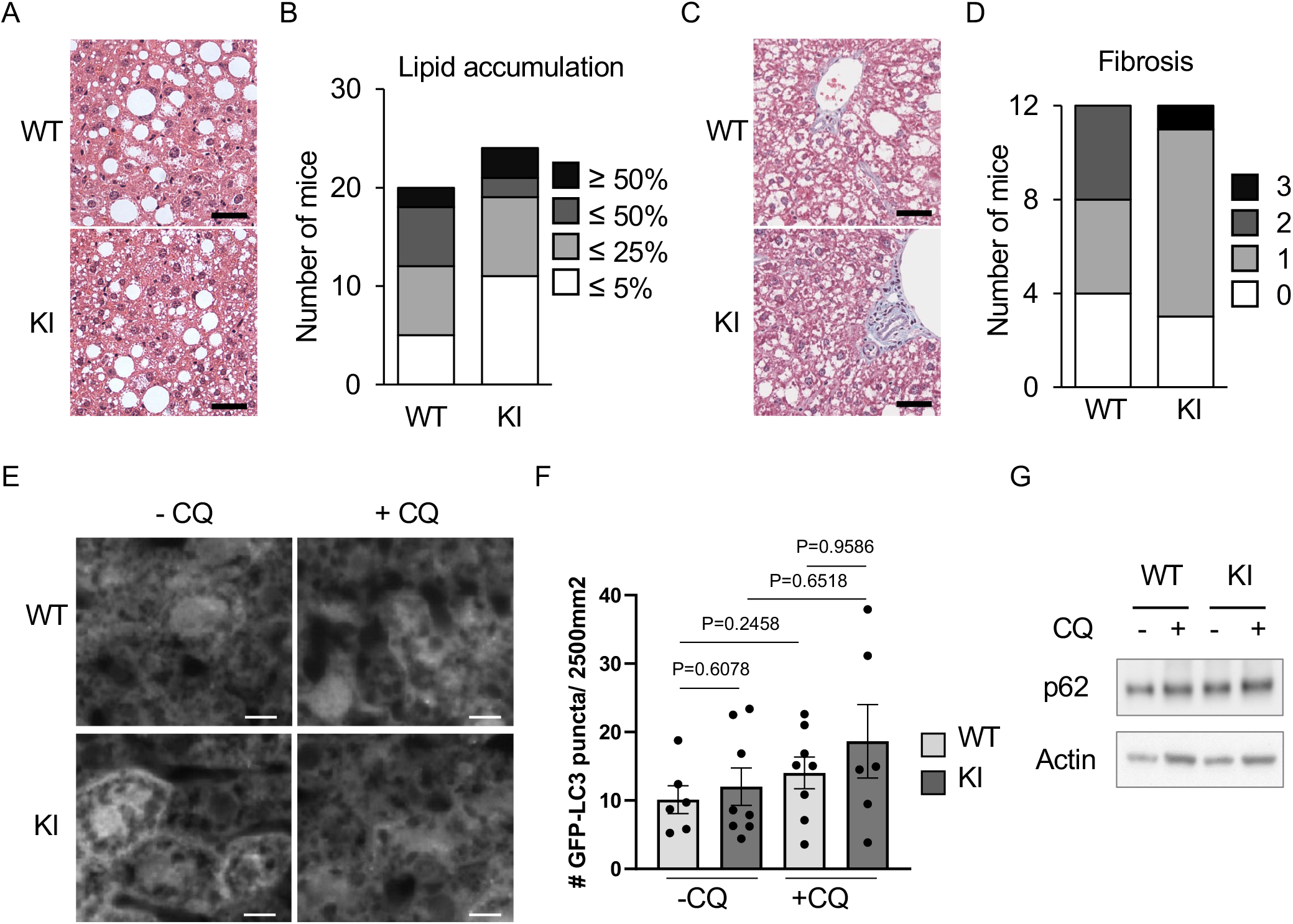
BECN1^F121A^ mutation does not improved liver age-related phenotype and does not increase autophagic flux in old mice. (**A**) Representative images of H&E stained liver sections of old *Becn1*^+/+^ (WT) and *Becn1^F121A/F121A^* (KI) mice. Scale bars, 40 μm. (**B**) Distribution of old mice according to the percentage of liver section area covered by lipid droplets (n=19 for WT, n=24 for KI). (**C**) Representative images of old WT and KI mice liver sections stained with Masson trichrome to quantify fibrosis visualized by the presence of collagen in light blue color. Scale bars, 40 μm. (**D**) Distribution of old mice according to their liver fibrosis score. (n=12 for WT and KI) (**E**) Representative images of GFP-LC3 puncta indicatives of autophagosomes in the liver of old WT and KI mice that transgenically express GFP-LC3, with or without chloroquine (CQ) for 6 h. Scale bars, 10 μm. (**F**) Quantification of GFP-LC3 puncta with or without CQ in old WT and KI mice. Data are mean ± s.e.m. (n=6 for WT and n=8 for KI without CQ and n=8 for WT and n=6 for KI with CQ). P values were determined by a two-sided unpaired t-test. (**G**) Western blot analysis of SQTM1/p62 autophagy marker and actin in the liver of old WT and KI mice. Shown are representative western blots of 3 independent experiments.

We next investigated whether in other organs, the increase in basal autophagy in *Becn1*^F121A^ KI mice was also impaired with age. As age-related cardiac and renal aging is prevented in the KI mice^28^, we measured autophagic flux in the heart and kidneys of old KI and WT mice expressing GFP-LC3. Compared to old WT mice, old KI mice had significantly more GFP-LC3 puncta in the heart (Fig. 2A and B), renal glomeruli (Fig. 2D and E) and renal proximal convoluted tubules (PCT) (Fig. 2F and G). We observed similar differences in GFP-LC3 puncta in the hearts and kidneys of old KI vs WT mice treated with CQ to block autophagic flux. Furthermore, in contrast to liver, CQ treatment further increased the number of puncta in both KI and WT mice, indicating that autophagic flux remains intact in the hearts and kidneys of 22 month-old mice. We also showed that the protein level of the autophagy substrate SQTM1/p62 is decreased in the heart and kidney of old KI mice compared to old WT mice (Fig. 2C and H). Thus, in contrast to liver, hearts and kidneys of old KI mice demonstrate greater autophagic flux compared to WT mice even at an advanced age.

**Figure 2.**
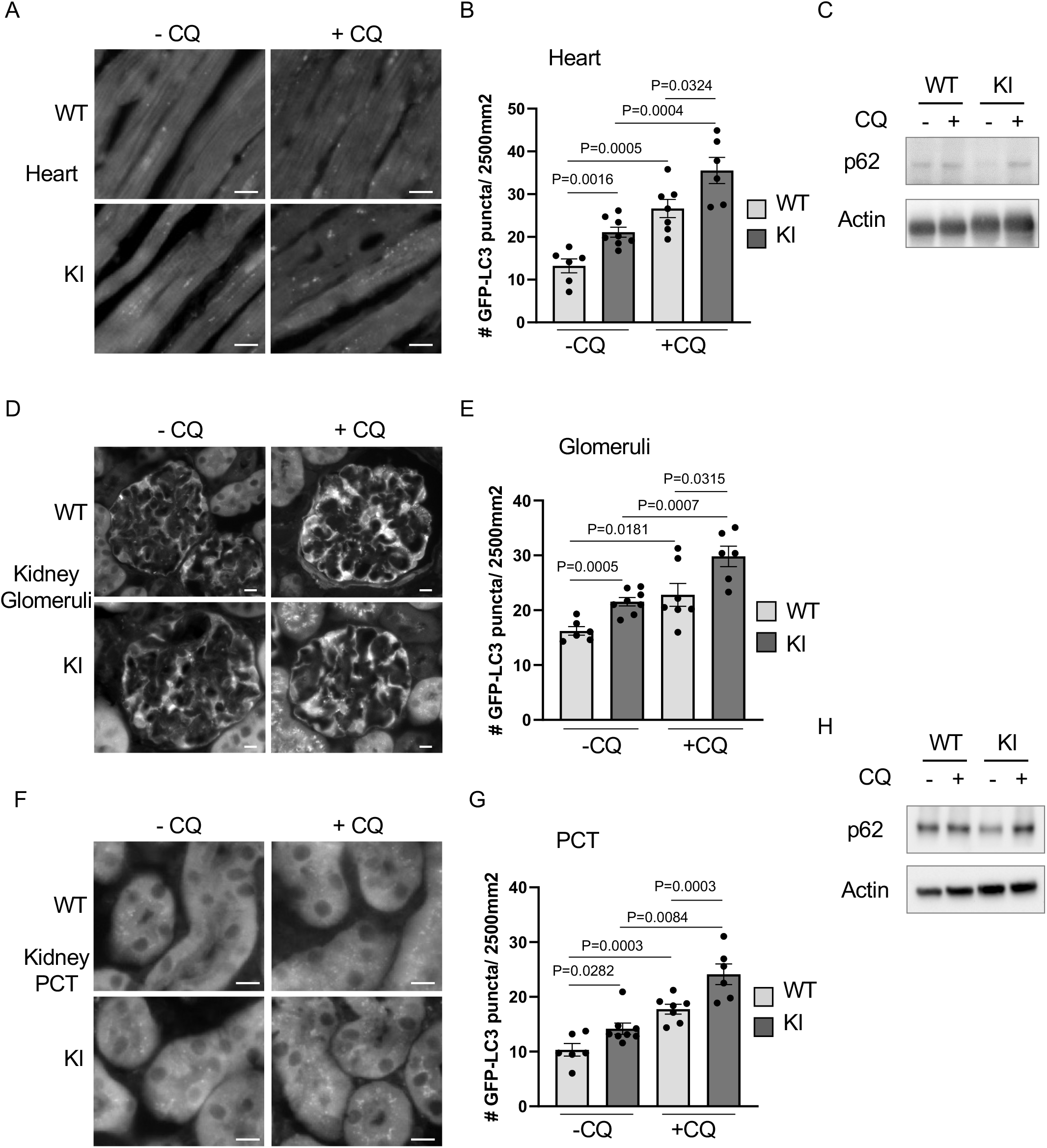
Autophagic flux is maintained in the heart and kidneys of old mice and is further increased by BECN1^F121A^ mutation. Representative images of GFP-LC3 puncta indicatives of autophagosomes in the heart (**A**), in the kidney’s glomeruli (**D**) and proximal convoluted tubules (PCT) (**G**) of old *Becn1* WT and KI mice that transgenically express GFP-LC3, with or without chloroquine (CQ) for 6 h. Scale bars, 10 μm. Quantification of GFP-LC3 puncta with or without CQ in the heart (**B**), in the kidney’s glomeruli (**E**) and PCT (**H**) of old *Becn1* WT and KI mice. Data are mean ± s.e.m. (n=6 for WT and n=8 for KI without CQ in all tissues and n=6 for KI and n=7-8 for WT with CQ). P values were determined by a two-sided unpaired t-test. Western blot analysis of SQTM1/p62 autophagy marker and actin, in the heart (**C**) and in the kidneys (**F**) of old *Becn1* WT and KI mice. Shown are representative western blots of 3 independent experiments.

To further investigate the potential of the *Becn1*^F121A^ KI mutation to increase autophagy during aging, we next focused on skeletal muscle. Using KI and WT mice expressing GFP-LC3, we observed that old KI mice had a statistically significant increased number of GFP-LC3 puncta in skeletal muscle compared to WT mice (Fig. 3A and B). Old KI mice treated with CQ also had a higher number of GFP-LC3 puncta than WT mice, indicating that autophagic flux is increased in skeletal muscle of old KI mice. It is worth noting that, like both heart and kidney, autophagic flux in the skeletal muscle of old mice remains intact as both WT and KI old mice treated with CQ had more GFP-LC3 puncta than untreated mice (Fig. 3B). The expression level of SQTM1/p62 is also lower in the muscle of old KI mice not treated with CQ compared to WT (Fig 3C). Altogether, these data indicate that old KI mice have increased autophagic flux in skeletal muscle.

**Figure 3.**
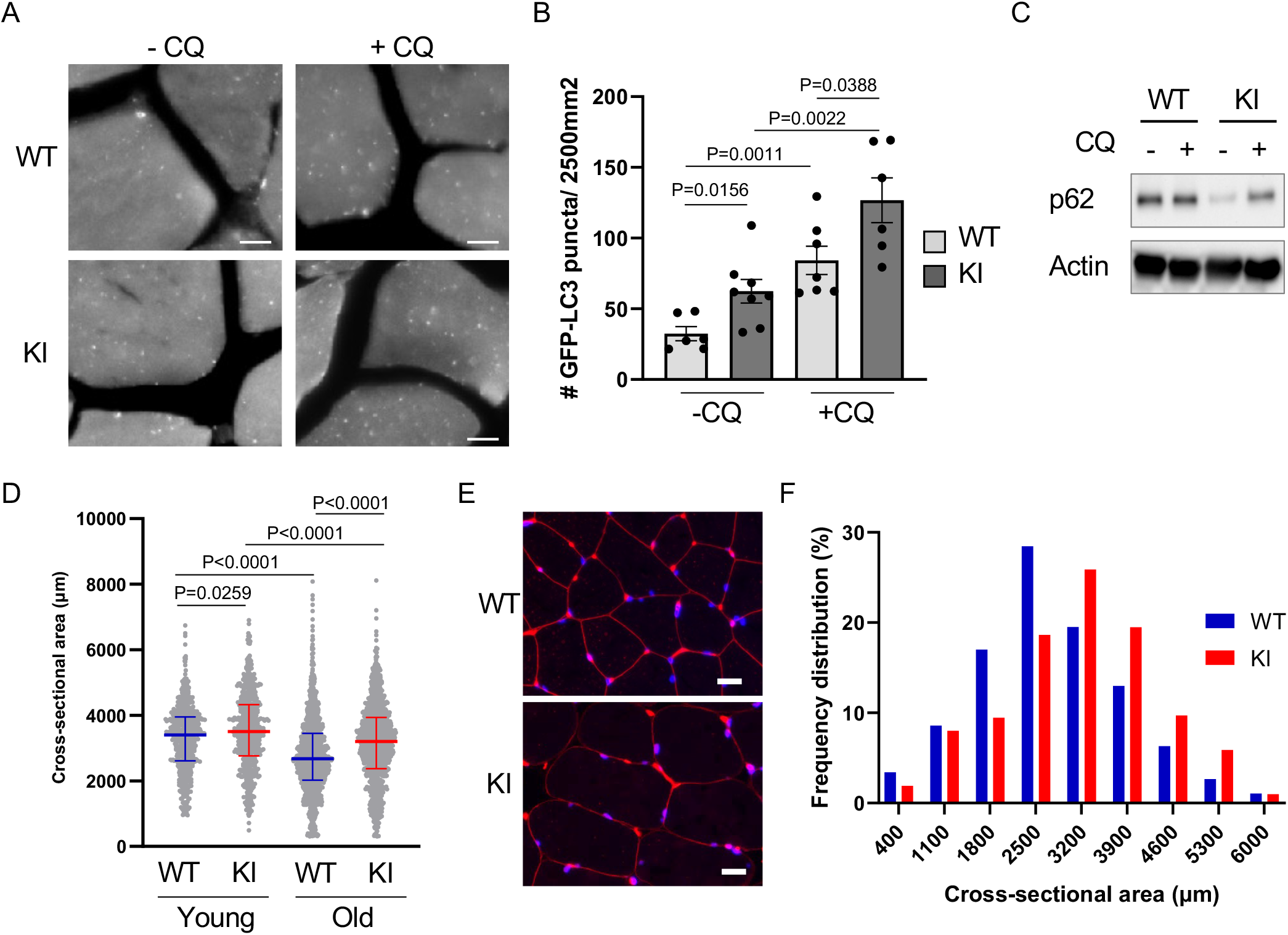
BECN1^F121A^ mutation increases autophagic flux in the skeletal muscle of old mice and prevents age-related decrease muscle fiber size. (**A**) Representative images of GFP-LC3 puncta indicatives of autophagosomes in the vastus lateralis of old *Becn1* WT and KI mice that transgenically express GFP–LC3, with or without chloroquine (CQ) for 6 h. (**B**) Quantification of GFP–LC3 puncta with or without CQ in old WT and KI mice. Data are mean ± s.e.m. (n=6 for WT and n=8 for KI without CQ and n=7 for WT and n=6 for KI with CQ). (**C**) Western blot analysis of SQTM1/p62 autophagy marker and actin, in the liver of old *Becn1* WT and KI mice. Shown are representative western blots of 3 independent experiments. (**D**) Cross-sectional area (CSA) of skeletal muscle fibers of the vastus lateralis muscle of young (5 month-old) and old (20 month-old) WT and KI mice (n=3 per group for young mice and n=5 per group for old mice). Graphs represents median CSA and interquartile and all myofiber CSA values are shown. (**E**) Representative images of vastus lateralis skeletal muscle sections of old WT and KI mice stained with a laminin antibody to outline the myofibers and DAPI. (**F**) Frequency of distribution of old WT and KI mice muscle fibers CSA (n=5 mice per group). Data represented as histograms of fiber size per CSA bin with 700 μm^2^ width. Scale bars, 10 μm. P values were determined by a two-tailed ANOVA with correction for multiple comparisons.

As increased autophagic flux in the heart and kidneys of old KI mice (Fig. 2) correlates with improved cardiac and renal aging phenotypes ^28^, we next investigated if old KI mice also had improved skeletal muscle aging phenotypes. One characteristic age-related change in the skeletal muscle is myofiber atrophy^35,36^. Indeed, when we measured the cross-sectional area of skeletal muscle fibers in young and old mice, we observed a clear age-related decrease in the muscle fiber size in WT mice (Fig. 3D and E). While myofiber size decreased with age in KI mice as well, the impact of aging was much less than in WT mice (Fig. 3D). Median myofiber cross-sectional area was significantly higher in old KI compared to old WT mice, although median myofiber cross-sectional area was similar between young KI and WT mice (Fig. 3D). Another characteristic of muscle aging is an increased heterogeneity in myofiber size, and this phenotype was more evident in old WT than in old KI mice. When we analyzed the frequency distribution of cross-sectional area of myofibers, we observed a clear shift towards larger fiber sizes in old KI compared to WT mice (Fig. 3F) suggesting that KI mice were protected from age-related myofiber atrophy. Thus, we conclude that old KI mice have increased autophagic flux and delayed muscle aging compared to old WT mice.

## Discussion

Our study indicates that in old mice, *Becn1*^F121A^ KI mutation increases autophagic flux in a tissue-specific manner and reduces aging phenotypes in the corresponding tissues/organs. Disruption of BECN1 binding to BCL2 by the KI mutation results in constitutively increased autophagic flux in all young mouse tissues explored to date: heart, kidneys, liver, skeletal muscles, mammary gland, adipose tissue, pancreas and brain^28,29,37,38^. However, as lysosomal function and autophagy activity have been described to decline with age, autophagic flux might also be inhibited. As a result, measuring autophagy by direct quantification of autophagy markers such as SQTM1/p62 and LC3 by immunoblot or fluorescence imaging under basal conditions may not properly reflect the autophagy level in aged animals and tissues^3,4^. Here, we investigated if autophagic flux remains increased by the *Becn*^F121A^ KI mutation in aged animals transgenically expressing GFP-LC3 and treated or not with the lysosomal inhibitor, chloroquine. Our data indicate that the constitutive increase in autophagic flux is maintained throughout aging in some tissues such as heart, kidneys and skeletal muscle but not in others like the liver, thus revealing an unexpected tissue-specificity in the upregulation of autophagy during mouse aging.

Our results also demonstrate that old WT mice maintain an active autophagic flux in the heart, kidneys and skeletal muscle and this autophagic flux can be increased further in old mice via expression of the BECN1^F121A^ mutant protein. These data also suggest that the concept that autophagic flux is repressed in old animals is not valid for all tissues, which is consistent with several studies that have also assessed autophagic flux in old mice using a lysosome inhibitor. Indeed, hematopoietic stem cells of old mice have an active basal autophagic flux that is even higher than in young mice^39^. Similarly, a recent study showed that autophagic flux increased with age in adipose tissue^40^. In line with our results, a study of autophagy in kidney aging found that old mice have higher autophagic flux than young mice in proximal convoluted tubules (PCT) but that further induction of autophagy in response to starvation only occurred in young mice^38^. Here we show that as opposed to starvation, the BECN1^F121A^ KI mutation can further increase autophagic flux in both PCT and glomeruli, which also maintain active autophagic flux during aging. Our results indicate that autophagy can be induced in old PCT and supports the hypothesis proposed by previously that the lack of autophagy response in old PCT in WT mice might be caused by dysregulation of the signaling pathways mediating autophagy induction in response to starvation rather than by lysosomal dysfunction preventing proper autophagy activity^38^. As the age-related renal phenotypes that are exacerbated in autophagy deficient mice are improved in old *Becn1^F121A^* KI mice, increasing autophagy could represent a potential strategy to alleviate age-related kidney diseases^28,38^. Autophagic flux is also increased in the hearts of old *Becn1^F121A^* KI mice, which is also consistent with a recent observation that exercise increases autophagic flux in old mice even though, as opposed to our result, this study did not detect any flux under basal resting condition^41^. This difference could be explained by the different assay used to measure flux. Although both studies used chloroquine to block lysosomal degradation, in the exercise study autophagy and flux were determined by immunoblot of LC3 and SQTM1/p62, whereas we used quantification of GFP-LC3 puncta combined with SQTM1/p62 analysis. Nevertheless, whether through genetic intervention via BECN1^F121A^ KI mutation, or a physiological intervention via exercise, both studies show that increased autophagic flux in the heart correlates with decreased cardiac aging in mice^28,41^. Similarly, our data show that in skeletal muscle, autophagic flux is not only active in aged mice but also higher in aged *Becn1^F121A^* KI mice and correlates with improved skeletal muscle aging. Previous studies have shown that autophagy deficiency in skeletal muscle leads to muscle loss and accumulation of protein aggregates which resemble accelerated muscle aging phenotypes in mice^11^. Likewise, recently caloric restriction was shown to improve skeletal muscle aging phenotypes and increased the number of autophagosomes, although flux was not assessed^42^. Our results demonstrate that increased autophagy in old *Becn1^F121A^* KI mice reduces age-associated skeletal muscle fiber atrophy, and provide supportive evidence for inducing autophagy as a potential therapeutic strategy to mitigate sarcopenia, the age-related decrease in skeletal muscle mass and strength.

In contrast to cardiac, renal and skeletal muscle aging, surprisingly hepatic aging was not improved in old *Becn1^F121A^* KI mice. Prior work had shown that autophagy deficiency in the liver leads to multiple dysfunctions such as hepatomegaly, mild injuries and the accumulation of abnormal organelles and lipid droplets in young mice^22,43,44^. In addition, deficiency in chaperone mediated autophagy accelerates liver aging^45^. The inability of the KI mutation to affect age-related liver damage phenotypes correlates with the absence of autophagic flux in old KI mice and absence of a difference between old KI and WT mice. As opposed to the heart, kidney and skeletal muscle, the constitutive increase in basal autophagy observed in young KI mice was not maintained with age in the liver of old KI mice. It is worth noting that the concept of an age-associated decline in autophagic flux was first established by the observation of decreased lysosomal activity and accumulation of autophagic and lysosomal vesicles with age in the livers of old mice and rats ^5,6,46^. Even if the BECN1^F121A^ KI mutation results in increased initiation and maturation of autophagosomes, downstream events such as reduced autophagosomal-lysosomal fusion and lysosomal degradation due to age-associated defects in their trafficking would still block autophagic flux and potentially explain why the BECN1^F121A^ KI mutation fails to increase autophagic flux in the livers of old mice^47^. In contrast to our findings, caloric restriction was shown to increase autophagic flux in the liver of old mice but only in female C57BL/6J mice^48^. Dysfunctions in autophagy flux have also been described in the aging brain^49^. However, our previous results indicate that age-related decline in autophagy is partially reversed in neural stem cells of 18 month-old *Becn1*^F121A^ KI mice^30^. Further investigation of autophagic flux in different cell populations of the brain as well as in other organs of old mice are required to better understanding the tissue-specificity of autophagy regulation during mammalian aging. Interestingly, though the BECN1^F121A^ KI mutation did not improve autophagic flux or the age-associated changes in the liver of old mice, the benefit of increased autophagy on other organs is sufficient to extends their lifespan^28^.

On a technical note, in our fluorescence imaging and analyses we observed an age-dependent accumulation of lipofuscin aggregates in all tissues, some of which appeared punctate. Accumulation of such aggregates is a well-established hallmark of aging and was especially pronounced in liver and PCT. As the lipofuscin fluorescence emission spectrum overlaps with GFP, puncate lipofuscin aggregates can potentially interfere with quantification of true GFP-LC3 puncta and as such should be carefully excluded.

To our knowledge, our study is the first to evaluate the autophagic flux in multiple tissues of old mice and highlights the importance of establishing a systemic evaluation of autophagic flux in aging mammals. We also demonstrate that increased autophagic flux in some old BECN1^F121A^ KI mouse tissues correlates with improved aging phenotypes in a tissue-specific manner. Overall, our data suggest that increasing autophagic flux during aging mitigates age-related phenotypes in multiple tissues.

## Materials and Methods

### Mice

*Becn1^F121A/F121A^* knock-in mice were generated in Beth Levine lab and backcrossed for more than 12 generations to C57BL/6J mice (Jackson Laboratories) as described ^29,28^. *Becn1^+/+^* (WT) and *Becn1^F121A/F121A^* (KI) littermate mice were crossed with GFP-LC3 transgenic C57BL/6J animals ^34^ and tissues of offspring were used for autophagic flux analyses. Old mice were 20 to 22 month-old littermates and young mice were 5 month-old. Both males and females were used for all analyses. All animal procedures were performed in accordance with institutional guidelines and with approval from the UT Southwestern Medical Center Institutional Animal Care and Use Committee.

### Autophagy analyses

To assess the autophagic flux in aged mice, 22-month-old homozygous *Becn1*^+/+^;GFP-LC3 or *Becn1^F121A/F121A^*;GFP-LC3 mice were synchronized by a 16 h starvation followed by 3 h of feeding before treatment with either PBS or chloroquine (50 mg kg^-1^) for 6 h. Mice were then perfused with 4% paraformaldehyde (PFA) in PBS and tissues were collected and processed for frozen sectioning as described ^28^. The mouse heart, liver, vastus lateralis skeletal muscle and kidney tissue sections were imaged using a 40× objective on a Zeiss AxioPlan 2 microscope. For each tissue, the total number of GFP-LC3 puncta was counted per 2,500 μm^2^ area (more than 20 randomly chosen fields were used per mouse) and was determined by an observer blinded to genotype. The average value for each tissue for each mouse was then calculated and graphed. For western blot analysis, tissues were lysed in ice-cold lysis buffer (Tris-HCl, pH 8, 300 mM, 2% SDS) with cOmplete, mini protease (Roche) and Halt phosphatase (Thermo Scientific) inhibitor cocktails for 30 min at 4 °C and the lysates were then centrifuged at 15,000*g* for 10 min. Cleared lysates were diluted in 2× SDS-PAGE loading buffer and submitted to western blotting using anti-p62 (GP62-C, Progen, 1:1,000 dilution), anti-LC3B (L7543, Sigma, 1:10,000 dilution), anti-BECN1 (sc-7382, Santa Cruz; 1:500 dilution), anti-BCL2 (sc-7382, Santa Cruz; 1:200 dilution) and anti-actin (sc-47778, Santa Cruz, 1:5,000 dilution) antibodies.

### Histopathological analyses

Mice were perfused with 4% PFA in PBS before tissue collection, fixation, and preparation of paraffin-embedded sections for histopathological analyses. Liver sections were stained with Hematoxylin and Eosin (H&E) then scanned using NanoZoomer 2.0-HT and analyzed using free NDPView2 software. To determine the lipid accumulation in the liver, each field of H&E stained liver sections was given a score using the following 4 categories: ≤5% tissue area; ≤25% tissue area; ≤50% tissue area; ≥50% tissue area with lipid droplets visualized as white empty vesicles on liver sections. For analyses of hepatic fibrosis, liver sections were stained with Masson’s trichrome according to the manufacturer’s instructions (ab150686, Abcam) and the sections were imaged using a 20× objective on a Zeiss AxioPlan 2 microscope. Ten random fields were evaluated per mouse and each field was given a fibrosis score using the following scale: 0, absence of damage; 1, ≤1% tissue area; 2, 1–5% tissue area; 3, ≥5% tissue area with fibrosis. The scores of each field were averaged to give a final fibrosis score for each mouse, ranging from 0 to 3. Quantification of all histopathological analyses was performed by an observer blinded to genotype.

### Muscle fiber size analyses

Skeletal muscle sections were staining with anti-laminin-2 (L0663, Sigma, 1:1000 dilution) to outline the muscles fibers. The average cross-sectional area of vastus lateralis muscles was determined using Myosight plugin ^50^ for FIJI (Just Image J) software. For old mice, 260 to 270 muscle fibers per mouse and 5 mice per genotype were analyzed. For young mice, 200 muscle fibers per mouse and 3 mice per genotype were analyzed.

### Statistical analyses

Data were analyzed using the GraphPad Prism 9 software. Two-tailed unpaired Student’s *t*-tests were used for analyses of autophagy. For the analysis of myofibers CSA, data were analyzed by two-way ANOVA with correction for multiple comparisons.

## Acknowledgments

We dedicate this article to the memory and legacy of Dr. Beth Levine whose intellectual and financial contributions were fundamental to this work. We thank Noboru Mizushima for the GFP-LC3 mice and Lori Nguyen for technical assistance. This work was supported by the Leducq Foundation grant 15CBD04 (S.S.) and NIH grant 5U19AI142784 (M.U.S.). The authors would also like to thank Linda W. and Milledge A. Hart III for their generous support of autophagy research.

## Abbreviations

CQ: chloroquine
GFP: green fluorescent protein
KI: BECN1^F121A^ knock-in mutation
MAP1LC3/LC3: microtubule associated protein 1 light chain 3
PCT: Renal proximal convoluted tubules

## Notes

### Competing Interest Statement

The authors have declared no competing interest.

## References

1. Mizushima N, Levine B. Autophagy in Human Diseases. N Engl J Med 2020; 383:1564–76.

2. Levine B, Kroemer G. Biological Functions of Autophagy Genes: A Disease Perspective. Cell 2019; 176:11–42.

3. Hansen M, Rubinsztein DC, Walker DW. Autophagy as a promoter of longevity: insights from model organisms. Nat Rev Mol Cell Biol 2018; 19:579–93.

4. Sarkis GJ, Ashcom JD, Hawdon JM, Jacobson LA. Decline in protease activities with age in the nematode Caenorhabditis elegans. Mech Ageing Dev 1988; 45:191–201.

5. Donati A, Cavallini G, Paradiso C, Vittorini S, Pollera M, Gori Z, Bergamini E. Age-related changes in the regulation of autophagic proteolysis in rat isolated hepatocytes. J Gerontol A Biol Sci Med Sci 2001; 56:B288–293.

6. Terman A. The effect of age on formation and elimination of autophagic vacuoles in mouse hepatocytes. Gerontology 1995; 41 Suppl 2:319–26.

7. Simonsen A, Cumming RC, Brech A, Isakson P, Finley DRS and KD. Promoting basal levels of autophagy in the nervous system enhances longevity and oxidant resistance in adult Drosophila. Autophagy 2007; 4:176–84.

8. Demontis F, Perrimon N. FOXO/4E-BP signaling in Drosophila muscles regulates organism-wide proteostasis during aging. Cell 2010; 143:813–25.

9. Bai H, Kang P, Hernandez AM, Tatar M. Activin signaling targeted by insulin/dFOXO regulates aging and muscle proteostasis in Drosophila. PLoS Genet 2013; 9:e1003941.

10. Kaushik S, Arias E, Kwon H, Lopez NM, Athonvarangkul D, Sahu S, Schwartz GJ, Pessin JE, Singh R. Loss of autophagy in hypothalamic POMC neurons impairs lipolysis. EMBO Rep 2012; 13:258–65.

11. Carnio S, LoVerso F, Baraibar MA, Longa E, Khan MM, Maffei M, Reischl M, Canepari M, Loefler S, Kern H, et al. Autophagy impairment in muscle induces neuromuscular junction degeneration and precocious aging. Cell Rep 2014; 8:1509–21.

12. Leidal AM, Levine B, Debnath J. Autophagy and the cell biology of age-related disease. Nat Cell Biol 2018; 20:1338–48.

13. Vinatier C, Domínguez E, Guicheux J, Caramés B. Role of the Inflammation-Autophagy-Senescence Integrative Network in Osteoarthritis. Front Physiol [Internet] 2018 [cited 2020 Nov 16]; 9. Available from: https://www.ncbi.nlm.nih.gov/pmc/articles/PMC6026810/

14. Meléndez A, Tallóczy Z, Seaman M, Eskelinen E-L, Hall DH, Levine B. Autophagy genes are essential for dauer development and life-span extension in C. elegans. Science 2003; 301:1387–91.

15. Hansen M, Chandra A, Mitic LL, Onken B, Driscoll M, Kenyon C. A Role for Autophagy in the Extension of Lifespan by Dietary Restriction in C. elegans. PLoS Genet [Internet] 2008 [cited 2020 Oct 19]; 4. Available from: https://www.ncbi.nlm.nih.gov/pmc/articles/PMC2242811/

16. Ulgherait M, Rana A, Rera M, Graniel J, Walker DW. AMPK modulates tissue and organismal aging in a non-cell-autonomous manner. Cell Rep 2014; 8:1767–80.

17. Kuma A, Hatano M, Matsui M, Yamamoto A, Nakaya H, Yoshimori T, Ohsumi Y, Tokuhisa T, Mizushima N. The role of autophagy during the early neonatal starvation period. Nature 2004; 432:1032–6.

18. Karsli-Uzunbas G, Guo JY, Price S, Teng X, Laddha SV, Khor S, Kalaany NY, Jacks T, Chan CS, Rabinowitz JD, et al. Autophagy is required for glucose homeostasis and lung tumor maintenance. Cancer Discov 2014; 4:914–27.

19. Ho TT, Warr MR, Adelman ER, Lansinger OM, Flach J, Verovskaya EV, Figueroa ME, Passegué E. Autophagy maintains the metabolism and function of young and old stem cells. Nature 2017; 543:205–10.

20. Sato S, Uchihara T, Fukuda T, Noda S, Kondo H, Saiki S, Komatsu M, Uchiyama Y, Tanaka K, Hattori N. Loss of autophagy in dopaminergic neurons causes Lewy pathology and motor dysfunction in aged mice. Sci Rep 2018; 8:2813.

21. Hara T, Nakamura K, Matsui M, Yamamoto A, Nakahara Y, Suzuki-Migishima R, Yokoyama M, Mishima K, Saito I, Okano H, et al. Suppression of basal autophagy in neural cells causes neurodegenerative disease in mice. Nature 2006; 441:885–9.

22. Komatsu M, Waguri S, Ueno T, Iwata J, Murata S, Tanida I, Ezaki J, Mizushima N, Ohsumi Y, Uchiyama Y, et al. Impairment of starvation-induced and constitutive autophagy in Atg7-deficient mice. J Cell Biol 2005; 169:425–34.

23. Komatsu M, Waguri S, Chiba T, Murata S, Iwata J, Tanida I, Ueno T, Koike M, Uchiyama Y, Kominami E, et al. Loss of autophagy in the central nervous system causes neurodegeneration in mice. Nature 2006; 441:880–4.

24. Rubinsztein DC, Mariño G, Kroemer G. Autophagy and Aging. Cell 2011; 146:682–95.

25. Pattingre S, Tassa A, Qu X, Garuti R, Liang XH, Mizushima N, Packer M, Schneider MD, Levine B. Bcl-2 antiapoptotic proteins inhibit Beclin 1-dependent autophagy. Cell 2005; 122:927–39.

26. Wei Y, Pattingre S, Sinha S, Bassik M, Levine B. JNK1-mediated phosphorylation of Bcl-2 regulates starvation-induced autophagy. Mol Cell 2008; 30:678–88.

27. Sinha S, Colbert CL, Becker N, Wei Y, Levine B. Molecular basis of the regulation of Beclin 1-dependent autophagy by the gamma-herpesvirus 68 Bcl-2 homolog M11. Autophagy 2008; 4:989–97.

28. Fernández ÁF, Sebti S, Wei Y, Zou Z, Shi M, McMillan KL, He C, Ting T, Liu Y, Chiang W-C, et al. Disruption of the beclin 1-BCL2 autophagy regulatory complex promotes longevity in mice. Nature 2018; 558:136–40.

29. Rocchi A, Yamamoto S, Ting T, Fan Y, Sadleir K, Wang Y, Zhang W, Huang S, Levine B, Vassar R, et al. A Becn1 mutation mediates hyperactive autophagic sequestration of amyloid oligomers and improved cognition in Alzheimer’s disease. PLoS Genet 2017; 13:e1006962.

30. Wang C, Haas M, Yeo SK, Sebti S, Fernández ÁF, Zou Z, Levine B, Guan J-L. Enhanced autophagy in Becn1F121A/F121A knockin mice counteracts aging-related neural stem cell exhaustion and dysfunction. Autophagy 2021;:1–14.

31. Chang JT, Kumsta C, Hellman AB, Adams LM, Hansen M. Spatiotemporal regulation of autophagy during Caenorhabditis elegans aging. eLife 2017; 6:e18459.

32. Kaushik S, Tasset I, Arias E, Pampliega O, Wong E, Martinez-Vicente M, Cuervo AM. Autophagy and the Hallmarks of Aging. Ageing Res Rev 2021;:101468.

33. Klionsky DJ, Abdel-Aziz AK, Abdelfatah S, Abdellatif M, Abdoli A, Abel S, Abeliovich H, Abildgaard MH, Abudu YP, Acevedo-Arozena A, et al. Guidelines for the use and interpretation of assays for monitoring autophagy (4th edition)1. Autophagy 2021; 17:1–382.

34. Mizushima N, Yamamoto A, Matsui M, Yoshimori T, Ohsumi Y. In vivo analysis of autophagy in response to nutrient starvation using transgenic mice expressing a fluorescent autophagosome marker. Mol Biol Cell 2004; 15:1101–11.

35. Sakellariou GK, Pearson T, Lightfoot AP, Nye GA, Wells N, Giakoumaki II, Vasilaki A, Griffiths RD, Jackson MJ, McArdle A. Mitochondrial ROS regulate oxidative damage and mitophagy but not age-related muscle fiber atrophy. Sci Rep 2016; 6:33944.

36. Brown M, Hasser EM. Complexity of age-related change in skeletal muscle. J Gerontol A Biol Sci Med Sci 1996; 51:B117–123.

37. Vega-Rubín-de-Celis S, Zou Z, Fernández ÁF, Ci B, Kim M, Xiao G, Xie Y, Levine B. Increased autophagy blocks HER2-mediated breast tumorigenesis. Proc Natl Acad Sci [Internet] 2018 [cited 2021 Dec 31]; Available from: https://www.pnas.org/content/early/2018/03/27/1717800115

38. Yamamoto S, Kuramoto K, Wang N, Situ X, Priyadarshini M, Zhang W, Cordoba-Chacon J, Layden BT, He C. Autophagy Differentially Regulates Insulin Production and Insulin Sensitivity. Cell Rep 2018; 23:3286–99.

39. Warr MR, Binnewies M, Flach J, Reynaud D, Garg T, Malhotra R, Debnath J, Passegué E. FOXO3A directs a protective autophagy program in haematopoietic stem cells. Nature 2013; 494:323–7.

40. Yamamuro T, Kawabata T, Fukuhara A, Saita S, Nakamura S, Takeshita H, Fujiwara M, Enokidani Y, Yoshida G, Tabata K, et al. Age-dependent loss of adipose Rubicon promotes metabolic disorders via excess autophagy. Nat Commun 2020; 11:4150.

41. Cho JM, Park S-K, Ghosh R, Ly K, Ramous C, Thompson L, Hansen M, Mattera MS de LC, Pires KM, Ferhat M, et al. Late-in-life treadmill training rejuvenates autophagy, protein aggregate clearance, and function in mouse hearts. Aging Cell 2021; 20:e13467.

42. Gutiérrez-Casado E, Khraiwesh H, López-Domínguez JA, Montero-Guisado J, López-Lluch G, Navas P, de Cabo R, Ramsey JJ, González-Reyes JA, Villalba JM. The Impact of Aging, Calorie Restriction and Dietary Fat on Autophagy Markers and Mitochondrial Ultrastructure and Dynamics in Mouse Skeletal Muscle. J Gerontol A Biol Sci Med Sci 2019; 74:760–9.

43. Ni H-M, Boggess N, McGill MR, Lebofsky M, Borude P, Apte U, Jaeschke H, Ding W-X. Liver-specific loss of Atg5 causes persistent activation of Nrf2 and protects against acetaminophen-induced liver injury. Toxicol Sci Off J Soc Toxicol 2012; 127:438–50.

44. Singh R, Kaushik S, Wang Y, Xiang Y, Novak I, Komatsu M, Tanaka K, Cuervo AM, Czaja MJ. Autophagy regulates lipid metabolism. Nature 2009; 458:1131–5.

45. Schneider JL, Villarroya J, Diaz-Carretero A, Patel B, Urbanska AM, Thi MM, Villarroya F, Santambrogio L, Cuervo AM. Loss of hepatic chaperone-mediated autophagy accelerates proteostasis failure in aging. Aging Cell 2015; 14:249–64.

46. Cuervo AM, Dice JF. Age-related decline in chaperone-mediated autophagy. J Biol Chem 2000; 275:31505–13.

47. Bejarano E, Murray JW, Wang X, Pampliega O, Yin D, Patel B, Yuste A, Wolkoff AW, Cuervo AM. Defective recruitment of motor proteins to autophagic compartments contributes to autophagic failure in aging. Aging Cell 2018; 17:e12777.

48. Mitchell SJ, Madrigal-Matute J, Scheibye-Knudsen M, Fang E, Aon M, González-Reyes JA, Cortassa S, Kaushik S, Gonzalez-Freire M, Patel B, et al. Effects of Sex, Strain, and Energy Intake on Hallmarks of Aging in Mice. Cell Metab 2016; 23:1093–112.

49. Nieto-Torres JL, Hansen M. Macroautophagy and aging: The impact of cellular recycling on health and longevity. Mol Aspects Med 2021;: 101020.

50. Babcock LW, Hanna AD, Agha NH, Hamilton SL. MyoSight—semi-automated image analysis of skeletal muscle cross sections. Skelet Muscle 2020; 10:33.

